# OMAMO: orthology-based model organism selection

**DOI:** 10.1101/2021.11.04.467067

**Authors:** Alina Nicheperovich, Adrian M. Altenhoff, Christophe Dessimoz, Sina Majidian

## Abstract

**Summary:** The conservation of pathways and genes across species has allowed scientists to use non-human model organisms to gain a deeper understanding of human biology. However, the use of traditional model systems such as mice, rats, and zebrafish is costly, time-consuming and increasingly raises ethical concerns, which highlights the need to search for less complex model organisms. Existing tools only focus on the few well-studied model systems, most of which are complex animals. To address these issues, we have developed **O**rthologous **Ma**trix and **M**odel **O**rganisms, a software and a website that provide the user with the best simple organism for research into a biological process of interest based on orthologous relationships between the human and the species. The outputs provided by the database were supported by a systematic literature review.

**Availability and implementation:** https://omabrowser.org/omamo/, https://github.com/DessimozLab/omamo

## 1 Introduction

Model organisms are non-human species used in human biomedical research to study development, gene regulation, and other cellular processes because they are relatively fast-growing, inexpensive, and easy to manipulate. Most importantly, their use has been possible due to the evolutionary conservation of biological processes (Wangler *et al*., 2017). Fast-moving progress in comparative genomics has allowed scientists to identify these evolutionary relationships by inferring human orthologs, genes that have diverged due to speciation (Fitch, 1970). Since orthologous genes tend to be functionally conserved and have common gene expression patterns, they are a better basis for model organism selection than other subtypes of homologs, which tend to functionally diverge faster (Altenhoff *et al*., 2012; Zheng-Bradley *et al*., 2010).

Currently used model organisms range from bacteria to complex mammals. The scientific community, however, aims to reduce the use of animals in research due to ethical implications, opting to use less complex organisms where possible. Currently available databases include MARVVEL (Wang et *al*., 2019), the Alliance of Genome Resources portal (Alliance of Genome Resources Consortium, 2000), and MORPHIN (Hwang et al., 2014). They focus on five to nine ‘traditional’ model organisms, most of which are complex organisms like mouse, rat and zebrafish. Moreover, their scope is restricted to human disease-related research. The only unicellular organisms considered in these databases are fission and budding yeast, whilst abundance of unicellular species in nature and their unique features make it difficult to find other non-complex model organisms for a biological process of interest.

To address the challenges above, we created an orthology-based database tool OMAMO alongside a user-friendly website that helps to select the best non-complex model organism for a biological process. Because the majority of species in the database have not been considered as model systems in the past, OMAMO has the potential to extend the set of organisms used in human biomedical research.

## 2 Methods

OMAMO takes advantage of the OMA database of orthologous genes. For a given biological process, the output presents a list of potential model organisms ranked based on their orthologous relationships with human.

For each species, pyOMA library was used to extract human orthologs (Altenhoff *et al*., 2021). For each ortholog, pyOMA was used to retrieve Gene Ontology (GO) terms, which provide information about the gene product and can represent one of the following three aspects: molecular function, cellular component and biological process (Gene Ontology Consortium, 2021). Some GO terms are general (e.g. ‘cell division’), whilst others are more specific (‘G2/M transition of mitotic cycle’). To quantify specificity of a GO term, we used information content (IC) calculated as *−log(p)* where *p* is its empirical frequency in the UniProt database (Pesquita, 2017), hence more specific GO terms have a higher IC value. The IC values were used to calculate functional similarity for each orthologous pair (Supplementary Section 1).

Orthologous pairs with functional similarity of < 0.05 were discarded. This aims to reduce the number of orthologs that only share general GO terms in the output. Consequently, gene pairs from a given species were grouped according to the biological process GO term they share. To maintain sufficient specificity in functional similarity considered, only GO terms with the IC value of ≥ 5 were kept. Finally, for each biological process GO term, species were ranked based on a scoring system, which takes into account the number of orthologs relevant to the biological process and average functional similarity across the genes.

We developed a freely accessible website for OMA (Fig.1), with the source code publically available. Out of the 50 species currently present in OMAMO, 31 are unicellular eukaryotes and the rest are bacteria. We suggest at least one model organism for 4620 out of 28,923 available biological GO terms (Gene Ontology Consortium, 2021). Since OMAMO is integrated in the OMA, it will be updated alongside the browser, meaning that the set of organisms will continue to grow and the database will include the latest GO annotations.

**Fig.1.**
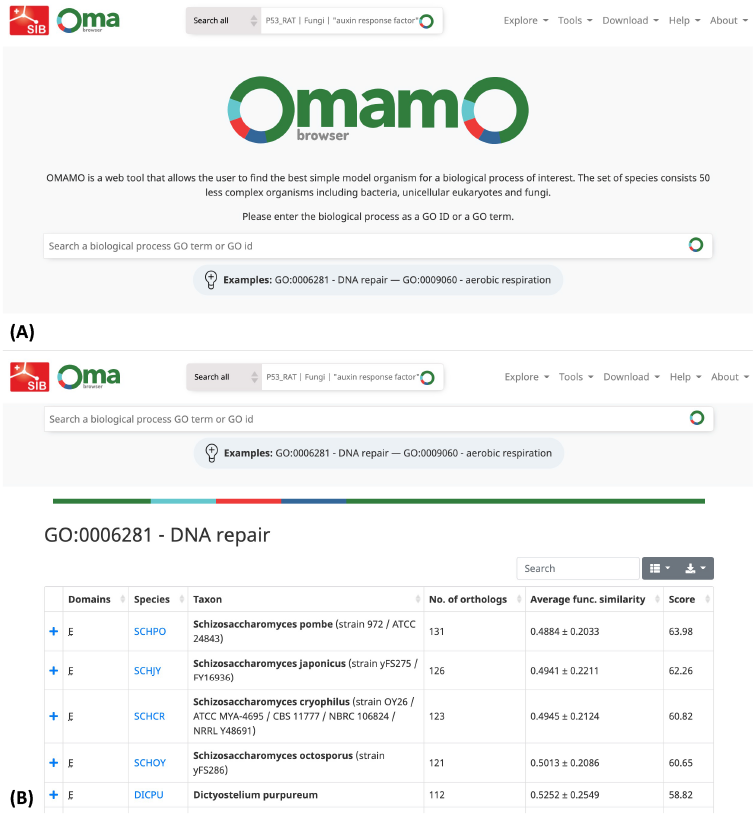
Website interface. (A) The main browser page of OMAMO. The user can search a GO term (‘DNA repair’) or a GO ID (0006281). (B) The output page gives a list of species ranked based on the score, but the user has the option to sort the output based on the total number of orthologs or the average functional similarity by clicking on the up-down sorting icon. The user can view orthologs by clicking on the ‘+’ button.

To validate our results, we referred to experimental evidence through a systematic literature search on PubMed (Supplementary Section 2). The top five review articles on three of the most well-studied organisms in OMA (*D. discoideum, N. crassa, S. pombe*) published in 2010-2021 were selected from the search output. Out of all biological processes which have been studied in one of the three organisms, the species of interest was in the top 5 model organism candidates in 42.6% of respective searches in OMAMO (Supplementary Section 2). This indicates that our algorithm is well supported by experimental data found in the literature.

## 3 Discussion

OMAMO is a freely-available database which aims to help scientists exploit alternative model species for human biomedical research. With the limited number of presently used model systems, the scientific community can now benefit from using other organisms, some of which could become model systems for processes that have previously only been studied in animals, leading to reduction in their use in experimental research. Moreover, this is the first database that provides such a wide range of potential model organisms. Due to the lack of literature on using species presented in OMAMO, the validation of results proved to be challenging. The following step for output validation would be to utilise proposed model species as model systems in wet-lab experiments. In the future, we plan to greatly expand the set of species and improve the scoring system by considering sequence similarity, conservation of protein structure and reproduction time. Additionally, we hope to provide unicellular model organisms for studying species other than human, for example animals for veterinary science research.

## Acknowledgements

The authors would like to thank Alex Warwick Vesztrocy and Natasha M. Glover for fruitful discussions.

## Supplementary Information

### 1. Method

#### 1.1 Functional Similarity Calculations

Functional similarity for each orthologous pair has been estimated using information content-based calculation of GO term overlap (Supplementary Equation 1), which has been shown to be a good predictor of functional relatedness between genes (Mistry and Pavlidis, 2008). GO terms have a hierarchical structure, i.e. general GO terms are at the top, whilst more specific (‘child’) terms are found in the lower branches of the GO hierarchy. The OMA browser provides only more specific GO annotations, which were then propagated to the parental terms up until a term with information content of 5 and below was reached. This was done to avoid domain terms (‘molecular function’, ‘cellular component’, and ‘biological function’) and other very general terms that are shared amongst almost all orthologs. Taking these terms into account would have led to a skewed distribution of functional similarity.

For two orthologous genes G_1_ and G_2_, we gather a set of GO terms for each ortholog, namely GO_1_ and GO_2_. Then, the functional similarity is defined using the following equations:

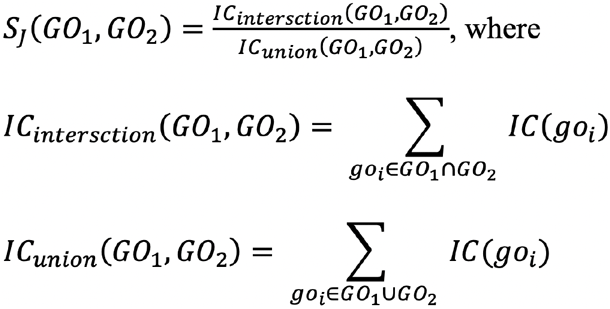

**Supplementary Equation 1.** Information content-based calculation of functional similarity (S_J_) of two orthologous genes based on Jaccard similarity (Popescu et al., 2006). It is measured as the ratio of information content (IC) of overlapping GO terms (and their parents) to the union of information content stored by GO terms of both genes (and their parental terms).

#### 1.2 Threshold settings

If the user chooses to use the website interface for their research, their result will be given with the following three filters set to default settings: (i) Minimal functional similarity of orthologous pairs (≥ 0.05) (Supplementary Figure 1); (ii) Minimum number of orthologous pairs per biological process (no threshold) (Supplementary Figure 2); (iii) Minimum IC value of GO terms (≥ 5) (Supplementary figure 3). However, if the user opts to use the software, they can change these settings according to their needs (e.g. if they wish to only consider orthologs with functional similarity of above 0.5 or if they only want to see species that have more than a certain number of orthologs for a given biological process).

**Supplementary Figure 1.**
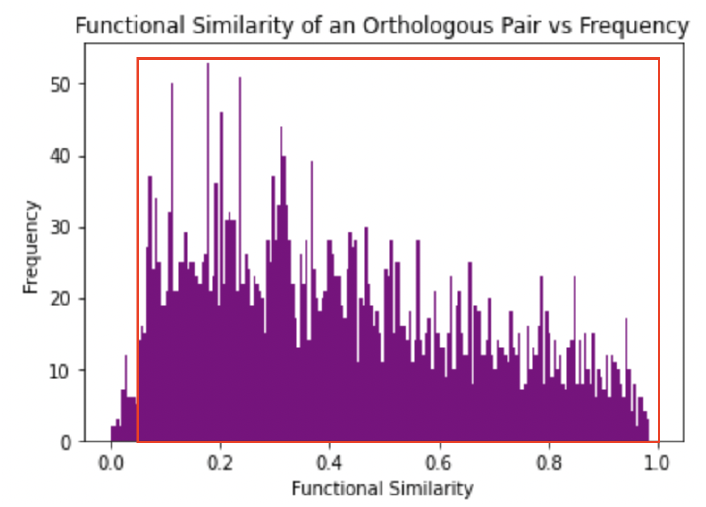
Distribution of functional similarity across orthologous pairs between *Dictyostelium discoideum* and *Homo Sapiens*. The red box shows default settings, i.e. the database only includes those orthologous pairs that have functional similarity of above 0.05. If the user wishes to only see outputs only for orthologs with high functional similarity, they can choose the threshold value to be higher.

**Supplementary Figure 2.**
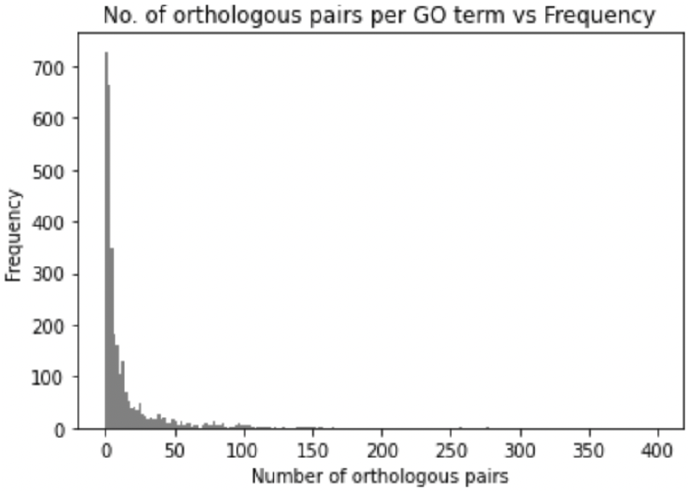
This histogram demonstrates distribution of orthologous pairs across biological process GO terms. For example, the peak to the left demonstrates that there are over 700 GO terms that have only one ortholog from *Dictyostelium discoideum* associated with them.

**Supplementary Figure 3.**
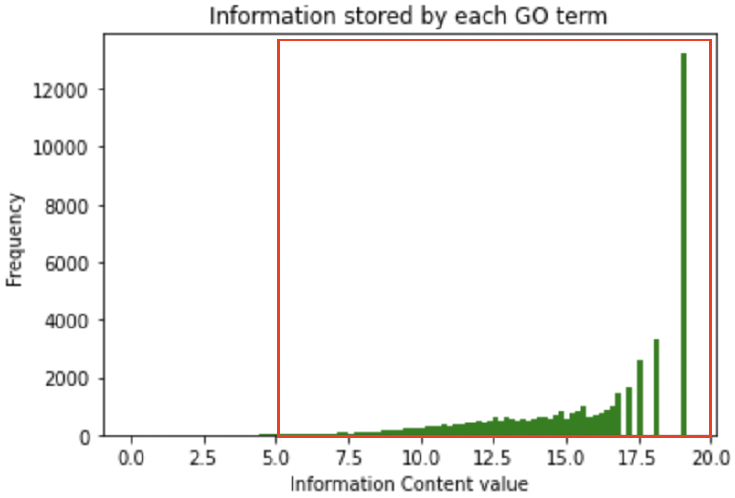
Distribution of Information content across all GO terms found in the UniProt as of July 2020. The red box shows the default setting of information content ≥ 5.

For example, if the user searches GO ID 0010737 (‘protein kinase A signalling’) with default settings, the output that they would get is like that shown in Supplementary Figure 4 (A). However, if the user changes the lower threshold for the number of orthologs in a model organism from 0 (default) to 4, their output would be different, as shown in Supplementary Figure 4 (B).

**Supplementary Figure 4.**
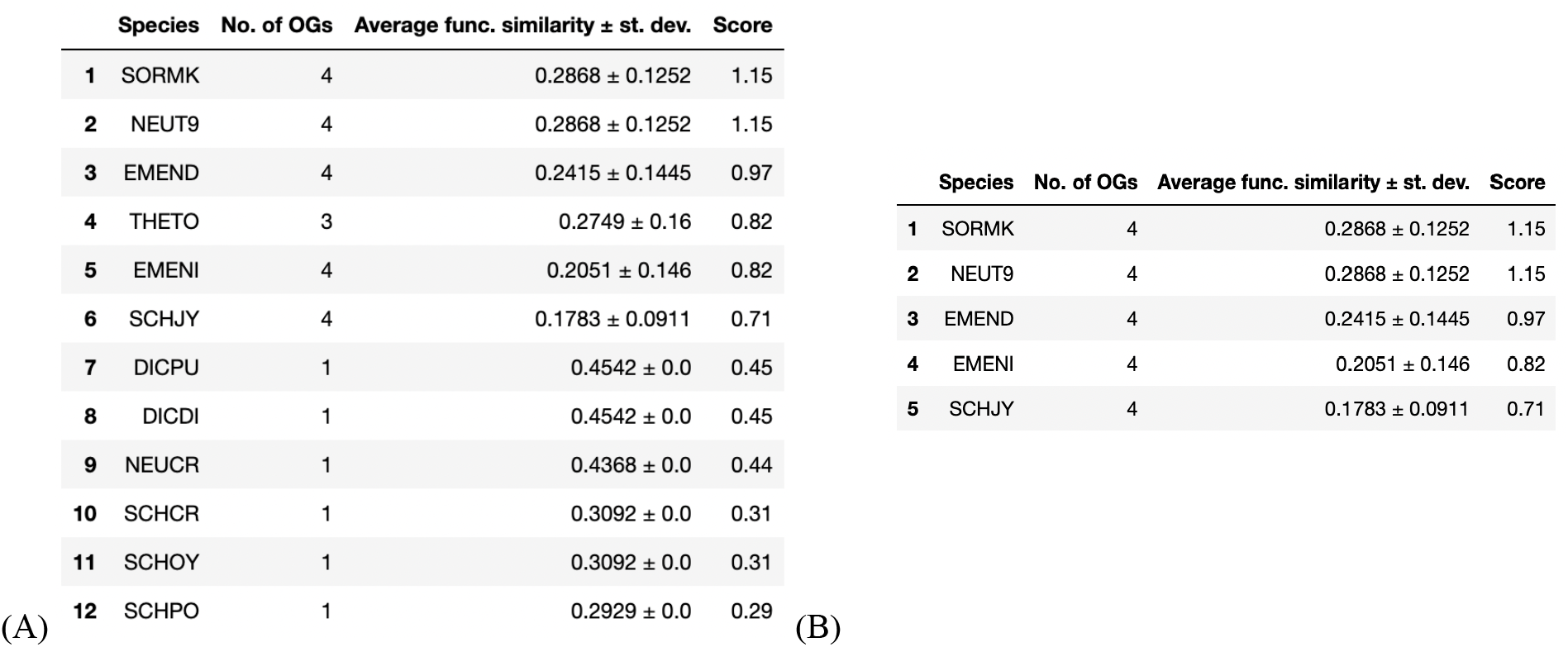
(A) Output for 0010737 with default settings. (B) Output for 0010737 where the lower threshold for number of orthologs has been set to 4.

Additionally, when using the code, the user can pick any combination of species present in OMA.

### 2. Systematic literature search

We chose 15 most relevant publications for the string ((species name) AND (model organism) AND (human)) in PubMed, five for each of the three species (*Dictyostelium discoideum, Neurospora crassa*, and *Schizosaccharomyces pombe*), from which we collected a total of 94 terms describing a biological process that has been studied in the species of interest. Searching for 21 of these gave no output or the species of interest wasn’t in the list of suggested model organisms, which is denoted as ‘-’ in the table below. In 40 out of 94 searches (42.6%), the species of interest was ranked as number 1-5. In 57 out of 94 searches (60.6%), the organism was in the top 10 suggested model systems, as summarised in the Supplementary Table 1 below.

**Supplementary Table 1.**

| Organism | Biological GO Terms (Rank No.) |
| --- | --- |
| ***Dictyostelium discoideum***<br>(Bozzaro, 2019; Pearce *et al.*, 2019; McLaren *et al.*, 2019; Martín-González *et al.*, 2021; Stuelten *et al.*, 2018) | Cell motility (6), chemotaxis (1), phagocytosis (1), macropinocytosis (-), cell adhesion (2), programmed cell death (4), autophagy (1), cytokinesis (14), lysosome organisation (2), cell-substrate adhesion (1), chemotaxis to cAMP (-), establishment of cell polarity (8), cell migration (1), phototaxis (-), mitochondrial transcription (1), protein insertion into mitochondrial outer membrane (-), aerobic respiration (2), mitochondrial organisation (10), mitochondrial localisation (11), thermotaxis(-), oxidative phosphorylation (14), reactive oxygen species biosynthetic process (1), mitochondrial calcium ion homeostasis (4), mitochondrial fission (19), macroautophagy (2), cellular water homeostasis (3), actin cytoskeleton organisation (2), protein phosphorylation (9), small GTPase-mediated signal transduction (2), multivesicular body assembly(-), multivesicular body organization (-), endosomal transport (2), vacuole organization (2), vacuolar transport (4), actin polymerisation and depolymerisation (2), regulation of cell-substrate adhesion (2), cellular component assembly (-), vesicle-mediated transport (3), intracellular signal transduction (3), regulation of signal transduction (11), regulation of intracellular signal transduction (3), protein processing (12), Notch receptor processing (-), amyloid precursor protein catabolic process (-), membrane protein intracellular domain proteolysis (1), ephrin receptor signalling pathway (-), organelle assembly (15), endosomal transport (2), multivesicular body assembly (-), energy derivation by oxidation of organic compounds (2), mitochondrial respiratory chain complex I assembly (6), mitochondrial electron transport NADH to ubiquinone (7), cellular protein complex assembly (15), mitochondrial ATP synthesis coupled electron transport (17), anion transport (4), regulation of biological quality (-). |
| ***Neurospora crassa***<br>(Jolma *et al.*, 2010; Ridenour *et al.*, 2020; Pelham *et al.*, 2020; Dunlap and Loros, 2017; Hevia *et al.*, 2016) | Circadian rhythm (-), histone methylation (19), histone H3 K27 methylation (-). |
| ***Schizosaccharomyces pombe***<br>(Allshire and Madhani, 2018; Matthews and Vosshall, 2020; Lin and Austriaco, 2014; Zhao, 2017; Florea, 2017) | Heterochromatin organisation (12), heterochromatin assembly (16), histone methylation H3K9 (-), gene silencing by RNA (5), cell division (2), programmed cell death(11), ageing (9), autophagy (9), necrotic cell death(-), intrinsic apoptotic signalling pathway (-), apoptotic process (6), regulation of cell death (4), mRNA splicing via spliceosome (6), RNA interference (8), mitochondrial inheritance (-), TOR signalling (4), response to reactive oxygen species (3), vacuolar acidification (10), mitochondrial fission (4), GMP biosynthesis (23), regulation of mitochondrial membrane potential (3), apoptotic DNA fragmentation (5), nuclear fragmentation involved in apoptotic nuclear change (-), reactive oxygen species process biosynthetic process (10), cell death (11), NAD<sup>+</sup> biosynthetic process (16), response to virus (1), regulation of cell cycle (5), regulation of vesicle-mediated transport (11), DNA replication (8), DNA repair (1), G2/M transition of mitotic cycle (2), mRNA processing (8), response to hydrogen peroxide (3), response to heat (8). |

